# Ageing and the ipsilateral M1 BOLD response: a connectivity study

**DOI:** 10.1101/2021.07.29.454012

**Authors:** Yae Won Tak, Ethan Knights, Richard Henson, Peter Zeidman

**Affiliations:** Wellcome Centre for Human Neuroimaging, University College London; MRC Cognition and Brain Sciences Unit, Department of Psychiatry, University of Cambridge

**Keywords:** ageing, aging, motor cortex, fMRI, DCM, connectivity, negative BOLD, ipsilateral

## Abstract

Young people exhibit a negative BOLD response in ipsilateral primary motor cortex (M1) when making unilateral movements, such as button presses. This negative BOLD response becomes more positive as people age. Here we investigated why this occurs, in terms of the underlying effective connectivity and haemodynamics. We applied dynamic causal modelling (DCM) to task fMRI data from 635 participants aged 18-88 from the Cam-CAN dataset, who performed a cued button pressing task with their right hand. We found that connectivity from contralateral supplementary motor area (SMA) and dorsal premotor cortex (PMd) to ipsilateral M1 became more positive with age, explaining 44% of the variability across people in ipsilateral M1 responses. Neurovascular and haemodynamic parameters in the model were not able to explain the age-related shift to positive BOLD. Our results add to a body of evidence implicating neural, rather than vascular factors as the predominant cause of negative BOLD – while emphasising the importance of inter-hemispheric connectivity. This study provides a foundation for investigating the clinical and lifestyle factors that determine the sign and amplitude of the M1 BOLD response, which could serve as a proxy for neural and vascular health, via the underlying neurovascular mechanisms.

## 1. Introduction

Making a unilateral movement is typically associated with activation of sensorimotor cortex in the contralateral hemisphere of the brain. However, such movements also engage the ipsilateral sensorimotor cortex [for recent reviews, see 1, 2]. This can result in a *negative* BOLD response (NBR) in the ipsilateral hemisphere, particularly in young people, as measured with fMRI. The ipsilateral NBR becomes less negative as people get older [3–6], in some cases becoming positive in sign with advanced age. This study investigates why ageing has this effect on the ipsilateral BOLD response.

The background to this question can be tackled into three parts: why is there a neural response to unilateral movements in ipsilateral sensorimotor cortex, why is the BOLD response negative for some people, and why does it become more positive with age?

Studies using EEG [7, 8] and ECoG [9] have demonstrated transient reductions in neural inhibition in both contralateral and ipsilateral sensorimotor cortex in response to unilateral movements. This is evidenced by decreases in alpha / mu and beta band spectral power (event-related desynchronization, ERD). Ipsilateral neural activity is sufficient to decode the precise movements being performed [10, 11], although it may not provide additional information beyond that encoded in the contralateral hemisphere [12]. What role does this ipsilateral neural activity play? A predominant view is that bilateral movements are the ‘default’ for the brain, with additional inter-hemispheric inhibition (IHI) required to enable unilateral movements and bimanual coordination, thereby suppressing mirror movements [13]. In support of this, most studies using transcranial magnetic stimulation (TMS) to target M1 have identified increased IHI during unilateral motor execution [1]. These studies typically apply a stimulus to motor cortex in one hemisphere, which reduces the motor-evoked potential generated by a subsequent stimulus applied to the opposite hemisphere [14]. This IHI is hypothesised to be mediated polysynaptically, with excitatory connections through the corpus callosum driving local GABA-ergic inhibition in the target hemisphere - as evidenced by the timing of TMS effects [15], intracranial recordings in animal models [16], pharmacology in humans [17] and modelling of neural dynamics [18]. With regards to behaviour, TMS studies have shown that disrupting ipsilateral motor cortex affects motor performance, for example, altering motor coordination and timing [19]. It should be noted, however, that there is considerable variability across TMS studies targeting ipsilateral motor cortex, with stimulation resulting in either increased or decreased IHI, and either facilitation or reduction of behavioural performance [1]. Nevertheless, the overall picture is that unilateral movements are associated with both hemispheres, albeit with less neural activity in the ipsilateral hemisphere. Nevertheless, the ipsilateral activity does carry information, and may have a functional role through reciprocal connectivity with the contralateral hemisphere.

Why is the BOLD response in ipsilateral sensorimotor cortex negative in sign, relative to rest, at least in young people? Unilateral medial nerve stimulation has been used to cleanly investigate the NBR. Stimulation causes a transient reduction in cerebral blood flow (CBF) and oxygen extraction (CMRO_2_) in ipsilateral sensorimotor cortex (S1/M1) [8], and similar results have been found for visual stimuli in occipital cortex [20]. With this in mind, to explain the generation of a NBR, one could simply invert the typical explanation for how a positive BOLD response (PBR) is formed. A reduction in neural activity due to inhibition would cause a decrease in CBF, outweighing a smaller decrease in CMRO_2_. This would increase the relative level of deoxyhaemoglobin (dHB), decreasing the BOLD signal relative to baseline. This explanation could be sufficient, were the NBR simply the inverse of the PBR. However, there is evidence that they differ in their neurovascular coupling. Higher metabolic demand (i.e., greater CMRO_2_/CBF coupling ratio) has been identified for NBRs relative to PBRs [8], perhaps caused by different neural cell types giving rise to them, each with differing metabolic demand. Thus, when identifying individual differences in the sign of the BOLD response, it is important to consider the potential for both neural and neurovascular contributions.

Having set out some key neural and haemodynamic determinants of the NBR, we next turn to ageing. Evidence from fMRI studies over the last 20 years has shown that older people display more positive BOLD responses in the ipsilateral sensorimotor cortex than younger people [3–5]. This effect co-locates with the ipsilateral NBR, which can be readily measured in younger, but not older, people [3]. Why might this change occur? There could be both neural and haemodynamic factors. At the neural level, inter-hemispheric connectivity changes with age. There is decline in the thickness of the corpus callosum [21] and a reduction in IHI from ipsilateral M1 to contralateral M1 [22]. This may underlie the observation that hemispheric asymmetries decrease with age, referred to as the Hemispheric Asymmetry Reduction in Older Adults (HAROLD) model [23]. An age-related shift in responses from M1 to higher level motor regions, such as dorsal premotor cortex (PMd), has also been noted. This “posterior to anterior shift in ageing” (PASA) [24] is supported by recent TMS results [22]. In parallel, ageing brings about changes in neurovascular coupling and haemodynamics. With advancing age, blood vessels stiffen, resulting in reduced cerebral blood flow dynamics (cerebrovascular reactivity) [25]. As set out above, NBRs have higher metabolic demand than PBRs, thus they could be particularly susceptible to any reduction in haemodynamic efficacy. To understand why the ipsilateral NBR changes with age, it is therefore important to distinguish neural and haemodynamic factors.

Our ultimate aim is to investigate how specific clinical and lifestyle factors impact on the ipsilateral NBR, which we hypothesise could serve as a marker for the health of the brain and its vasculature. In this foundational paper, we use dynamic causal modelling (DCM) of fMRI data in order to quantify neural and haemodynamic contributions to the NBR. Previously, DCM has been applied successfully to study the motor system in ageing, with results that are generally consistent with findings from TMS [26, 27]. Here, we applied this approach to quantify the evidence for three sets of hypotheses for why the NBR in ipsilateral M1 decreases with age. We asked to what extent age-related changes in NBR could be explained by 1) reorganisation of neural networks, particularly inter-hemispheric connections, 2) individual differences in the strength of neurovascular coupling, and / or 3) changes in the rate of blood flow.

To address this, we leveraged a unique BOLD fMRI dataset from the Cam-CAN project [28, 29], in which n=635 participants aged 18-88 performed a simple motor response task with their right hand, in response to visual and auditory cues. The data suggested two types of subject – those with mainly negative BOLD responses in right M1, who tended to be younger, and those with mainly positive right M1 responses, who tended to be older. We then specified biophysical models – i.e., dynamic causal models (DCMs) – which included neural, neurovascular and haemodynamic components. Each of these had parameters that we estimated from the fMRI data. We identified the parameters that could best explain differences in right M1 response across individuals of different ages, and we performed simulations to identify the minimal set of parameters that could explain the age-related shift from NBR to PBR.

## 2. Materials and Methods

### 2.1 Participants and task

We used openly available data from the Cambridge Centre for Ageing and Neuroscience (Cam-CAN) project [28, 29] (available at http://www.mrc-cbu.cam.ac.uk/datasets/camcan/). The participants had been screened to ensure no Magnetic Resonance (MR) contradictions, no learning disability, no cognitive impairment (tested through Mini-Mental State Examination with a score >24) and no long-term illness or disability. From an initial sample of 652 of participants with fMRI data, we excluded nine participants with missing data or who had fMRI signal dropout. We further excluded eight participants who had suffered a stroke, giving a final sample size of n=635.

While undergoing fMRI, participants responded to 128 trials of a simple audio/visual sensorimotor task. On each trial, a binaural auditory tone (300Hz, 600Hz or 1200Hz) was played and two round checkerboards were presented simultaneously to the left and right of a central fixation cross. Auditory tones had a duration of 300ms, and checkerboards had a duration of 34ms. 120 trials were presented (40 per auditory frequency). Additionally, there were four trials when only an auditory tone was played and four trials where only checkerboards were presented. Importantly, randomly interspersed null trials were included with stimulus onset asynchronies (SOAs) from 2s to 26s, enabling efficient estimation of the haemodynamic response. The participant’s task was simply to press a button with the right index finger when they saw or heard a stimulus.

A recent detailed analysis of the in-scanner behavioural outcomes of this task (in these same subjects) identified no effect of age on the mean reaction time (since the task was unspeeded), but there was a significant effect of age on the variability in reaction times across trials [12]. No significant relationship between performance and the BOLD response was found, therefore we did not model reaction times in the analyses that follow.

### 2.2 fMRI data and pre-processing

We used pre-processed fMRI data from the Cam-CAN project from the “aamod_norm_write_dartel” analysis stage. Details of scanning sequences and the image pre-processing pipeline can be found in Ref [29]. We applied an additional spatial smoothing, with an isotropic Gaussian kernel of 3mm at its full-width half maximum, which resulted in a final estimated smoothness of the group level maps of 9mm isotropic, which meets the assumptions of random field theory that the image smoothness is at least three times the voxel size (which was 3mm here).

### 2.3 SPM analysis

To identify brain regions responding to the task, we performed a preliminary Statistical Parametric Mapping (SPM) analysis using SPM version 12 (https://www.fil.ion.ucl.ac.uk/spm/). Each subject’s first level general linear model (GLM) included the three experimental conditions (Audio+Visual, Audio, Visual), and the 3 translations and 3 rotations from motion correction. Second level models (one sample t-tests) included covariates for the linear and non-linear effects of age (age, age squared and age cubed), in addition to potential confounds such as height, weight and hearing loss. Full details of the SPM analysis, including a table of confounding variables, are provided in the supplementary materials.

### 2.4 Region of interest definition

We defined binary masks for six regions of interest (ROIs). Right primary motor cortex (M1), right dorsal PMd and bilateral supplementary motor areas (SMA) were defined based on two criteria: 1) voxels were only included if they showed a significant effect of age on Auditory+Visual trials at p < 0.05 family-wise error corrected (as assessed by a group-level F-contrast over age, age squared and age cubed), and 2) the same voxels had to overlap with corresponding motor regions in the Human Motor Area Template (HMAT) atlas [30]. The remaining two regions, left M1 and left PMd, did not show age effects, and so were defined as the mirror image around the midline of right M1 and right PMd ROIs respectively. In what follows, we refer to the left (contralateral) hemisphere regions as lM1, lPMd and lSMA, and the right (ipsilateral) hemisphere regions as rM1, rPMd and rSMA.

### 2.5 Timeseries extraction

For each subject and ROI, we extracted a representative timeseries for subsequent DCM analyses. Following standard procedures in SPM, this was the first principal component across voxels within the ROI. To reduce noise, contributing voxels were limited to those exhibiting a response to the task at a liberal statistical threshold (p < 0.001 uncorrected) at the individual subject level. Where no voxels could be found at this threshold, the threshold was iteratively relaxed to p < 0.01, p < 0.1 and finally p < 1, until at least one voxel was included. Each representative timeseries was then high-pass filtered, pre-whitened and corrected for confounds (to remove the mean and head motion) in the usual way using the SPM software.

### 2.6 Dynamic causal model

We used the standard (one-state, deterministic) DCM for fMRI, with certain changes to the parameters for this study. We begin by briefly reprising the form of the model.

The inputs to the DCM were the fMRI timeseries data and experimental timing. Matrix ***Y***_***i***_ ∈ ℝ_*V×R*_ were fMRI timeseries for subject *i* = 1 ... *N*, with *V* = 261 measurements from *R* = 6 brain regions. Columns of the matrix ***U***_***i***_ ∈ {0,1}_*T×P*_ encoded the onsets of the *P* = 3 experimental conditions (auditory+visual, auditory only, visual only) over *T* = 4176 time bins (16 bins per TR).

The DCM for fMRI model is split into two parts: neural and haemodynamic. The neural part has one hidden state variable per brain region – encoding the level of neural activity ***z***_***i***_(*t*) ∈ ℝ_*R*_. The dynamics of this activity are governed by the differential equation:

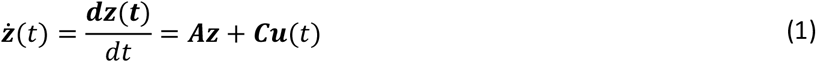

Where parameter matrix ***A***_*R×R*_ ∈ ***θ***_***i***_ is a connectivity or directed adjacency matrix and parameter matrix ***C***_*R×P*_ ∈ ***θ***_***i***_ encodes the strength of driving input due to each experimental condition to each brain region. Element *A*_*ij*_ is the rate of change of neural activity in region *j* due to region *i*, which is called the *effective connectivity* and has units of Hertz (Hz). To ensure stability, the elements of the leading diagonal of matrix ***A*** (self-connections) must be negative in sign. Therefore, rather than estimate those values directly, they are replaced with log scaling parameters 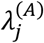, which are estimated from the data and are translated to units of Hz by the function:

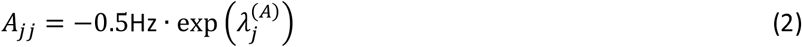

where the exponential ensures positivity of the scaling parameter, and −0.5Hz is the default strength of self-inhibition.

The haemodynamic part of the model translates neural activity to predicted fMRI timeseries. It adds hidden states encoding blood volume, deoxyhaemoglobin and cerebral blood flow. This part of the model has two types of free parameter. Rate parameter *κ* controls the rate of decay of the vasoactive signal (for example, nitric oxide), which couples neural activity to the haemodynamic response. It is estimated from the data via a log scaling parameter *λ*^(*κ*)^, which is translated to units of Hz via the definition *κ* = 0.64Hz · exp(*λ*^(*κ*)^). The remaining parameters are the haemodynamic transit times. These are time constants, *τ*_*j*_ for each region *j*. They quantify the rate of blood flow through the venous compartment. They are in units of seconds and are estimated from the data via the introduction of a log scaling parameter for each region 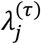, defined as 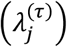. For the full definition of the haemodynamic model, please see Appendix 5 of [Ref. 31].

Given the importance of accurate haemodynamic modelling in this study, we made several updates to the generative model in DCM for fMRI model to optimize it for our 3T fMRI data. We set the echo time (TE) to use the precise value for this experiment (0.03s), rather than the fixed default value of 0.04s (in the Matlab function spm_gx_fmri.m). Table 1 lists the other parameters we adjusted for this study with details in the footnotes.

**Table 1.**
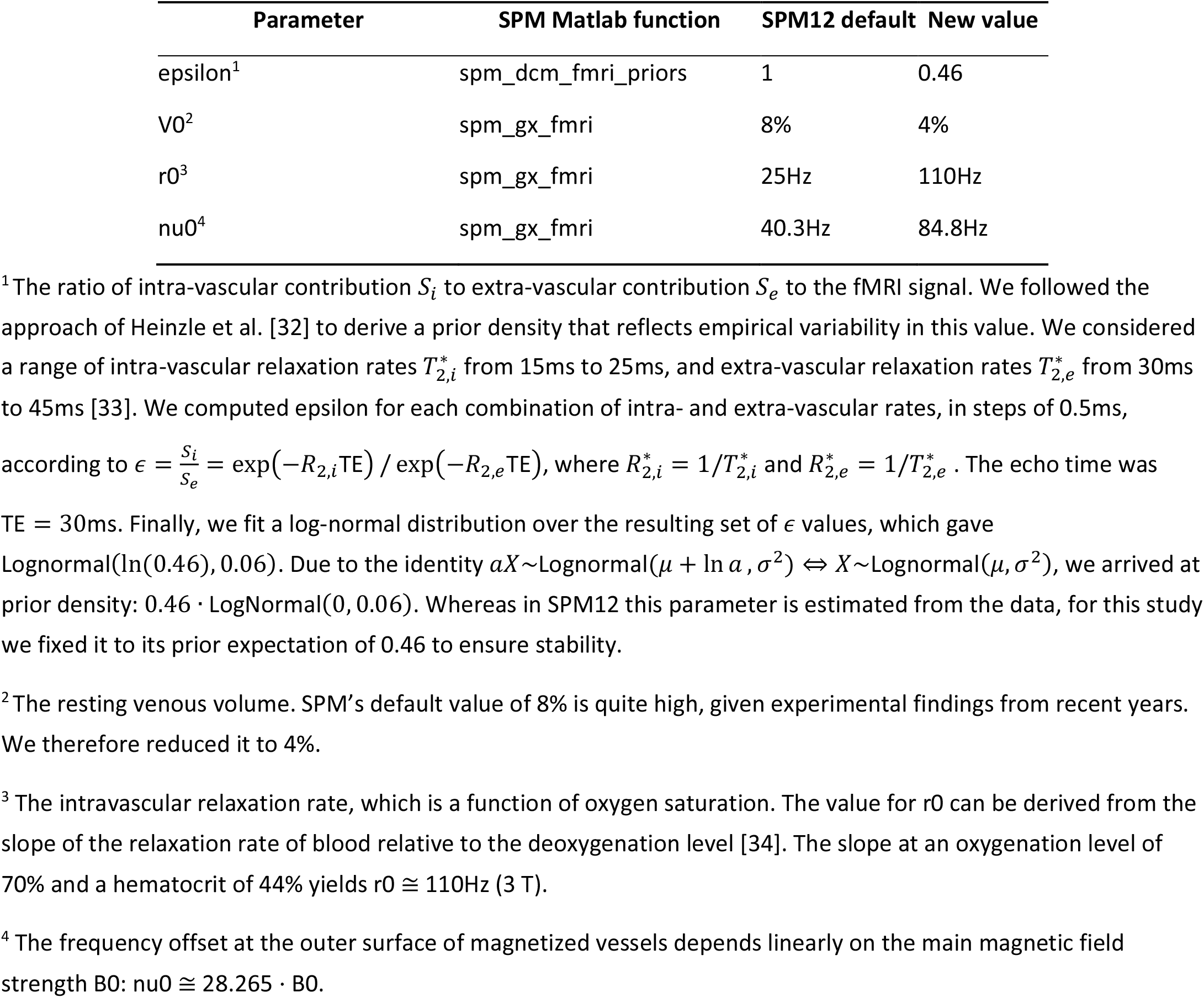
Updated DCM parameters

### 2.7 Priors on DCM parameters

Each DCM for subject *i* had free parameters ***θ***_***i***_ = {***A***, ***λ***^(*A*)^, ***C***, *λ*^(*κ*)^, ***λ***^(***τ***)^}, i.e., between-region connectivity, self-connections, driving input, decay rate and transit time. The other parameters listed in Table 1 were fixed values (they had infinite precision). We specified two DCMs per subject, *m*_1_ and *m*_2_, which differed in the location of the driving inputs (detailed in Figure 2). Parameters were switched on or off in each model through specification of the prior probability density *p*(***θ***_***i***_|*m*_1_) = *N*(***μ***_**1**_, **Σ**_**1**_) and *p*(***θ***_***i***_|*m*_2_) = *N*(***μ***_**2**_, **Σ**_**2**_) respectively. Parameters with prior variance close to zero – i.e., small values on the leading diagonal of prior covariance matrix **Σ**_**1**_ or **Σ**_**2**_ – were effectively ‘switched off’, fixing them at their prior expectation ***μ***_**1**_ or ***μ***_**2**_ respectively (typically zero). Whereas, parameters with larger positive prior variances were ‘switched on’ and informed by the data. For full details of the priors in DCM for fMRI, see Table 3 of [Ref. 31].

We assigned models *m*_1_ and *m*_2_ the same priors on their connectivity parameters (***A***, ***λ***^(*A*)^), such that all connections among the six regions were informed by the fMRI data (i.e., all connections were switched on). This was based on evidence for both homotopic and non-homotopic connections among motor cortex regions, from tract-tracing in non-human primates [35] and diffusion tensor imaging (DTI) in humans [36, 37]. (For a brief review of non-homotopic premotor connections, see [Ref. 22]).

We set priors on the driving inputs (parameter matrix ***C***) in model *m*_1_ such that lSMA and lPMd could receive driving input by the task, and in model *m*_2_ rSMA and rPMd could receive driving input. Any dynamics in M1 were therefore mediated by connections from SMA and PMd. This expressed the hypothesis that there is a motor hierarchy, with SMA and PMd at a higher level than M1. A similar architecture has been used in previous DCM analysis of the motor cortex [38].

### 2.8 Model estimation and PEB

We fitted the two DCMs per subject to the fMRI data using the standard Bayesian scheme in SPM (Variational Laplace [39]). To attempt to rescue any subjects whose estimates were stuck in local optima, we then re-estimated every model using the group average connectivity as the starting value of the estimation (while leaving the priors untouched). The final outputs of the model estimation procedure were the posterior probability densities over the parameters for each of the two models per subject, *p*(***θ***_***i***_|***Y***_***i***_, *m*_1_) and *p*(***θ***_***i***_|***Y***_***i***_, *m*_2_), and the log model evidences, approximated by the free energy 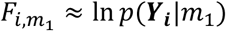 and 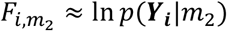. Each free energy scores the quality of the model, in terms of its accuracy minus its complexity, enabling Bayesian model comparison.

We then used the Parametric Empirical Bayes (PEB) scheme in the SPM software to identify commonalities and differences in parameters at the group level. The software stacked the *M* parameters from all *N* subjects into a vector 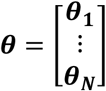 and modelled them at the group level using the general linear model:

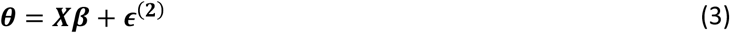

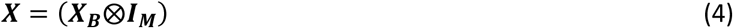

The between-subjects design matrix ***X***_*B*_ ∈ ℝ_*N*×4_ contained four orthogonal regressors: a constant (all ones), group (1 for negative responders, 0 for positive responders, before mean-centring), the residual effect of age after accounting for group, and the residual level of rM1 BOLD activity after accounting for group and age. The latter two regressors were ‘nuisance effects’, included to maximize the explained variance of the model, and were calculated using recursive Gram-Schmidt orthogonalisation implemented in the SPM Matlab function: spm_orth.m. This procedure also mean-centred the regressors in ***X***_*B*_, making the first regressor (the constant) interpretable as the group average connectivity. The Kronecker-tensor product ⨂ with the identity matrix ***I***_*M*_ was applied automatically by the software, to replicate the four between-subject effects over the *M* DCM parameters per subject. The resulting design matrix ***X*** = ℝ_*NM*×4*M*_ contained a regressor for each covariate on each DCM parameter. Finally, the between-subjects variability ***ϵ***^(**2**)^ was modelled using a covariance component model. We elected to have one component per field (A, C, decay, transit), allowing each type of parameter to have a different level of between-subjects variability.

We specified a PEB model for each of the two DCMs (left hemisphere driving and right hemisphere driving), and estimated each PEB model to obtain a posterior probability densities over the parameters, 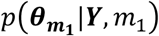 and 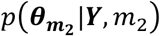, as well as the free energy for each hierarchical model 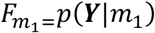 and 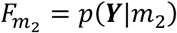, where ***Y*** was the complete dataset from all subjects.

### 2.9 Bayesian model comparison

We tested hypotheses by comparing the evidence for different models (Bayesian model comparison). This procedure identified the model(s) that offered the optimal trade-off between model accuracy and complexity. We began by comparing the evidence for the two PEB models described above, which differed in whether left or right hemisphere regions served as driving inputs, i.e. the log Bayes factor was 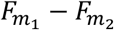. We then performed a series of comparisons of a full PEB model against reduced models where different mixtures of regressors (columns of the design matrix ***X*** in Eq. 3) were switched off, by setting restrictive priors on the corresponding regression parameters (***β***). The probability for each parameter being present versus absent was calculated by performing repeated Bayesian model comparisons, with each parameter switched on versus switched off.

## 3. Results

### 3.1 Participant demographics

Of 635 participants, 319 were male (50%), aged from 18 to 88 (18-28: n=50, 28-38: n=102, 38-48: n=99, 48-58: n=98, 58-68: n=100, 68-78: n=99, 78-88: n=87).

### 3.2 Region of interest (ROI) analyses

A preliminary mass-univariate (SPM) analysis identified significant effects of age in bilateral SMA (left: 52 voxels, right: 66 voxels) and right premotor/motor regions (rPMd: 38 voxels and rM1: 188 voxels). There were no significant effects of age in left premotor/motor regions (lPMd and lM1). We therefore specified three ROIs per hemisphere: SMA, PMd and M1 (Figure 1A).

**Figure 1.**
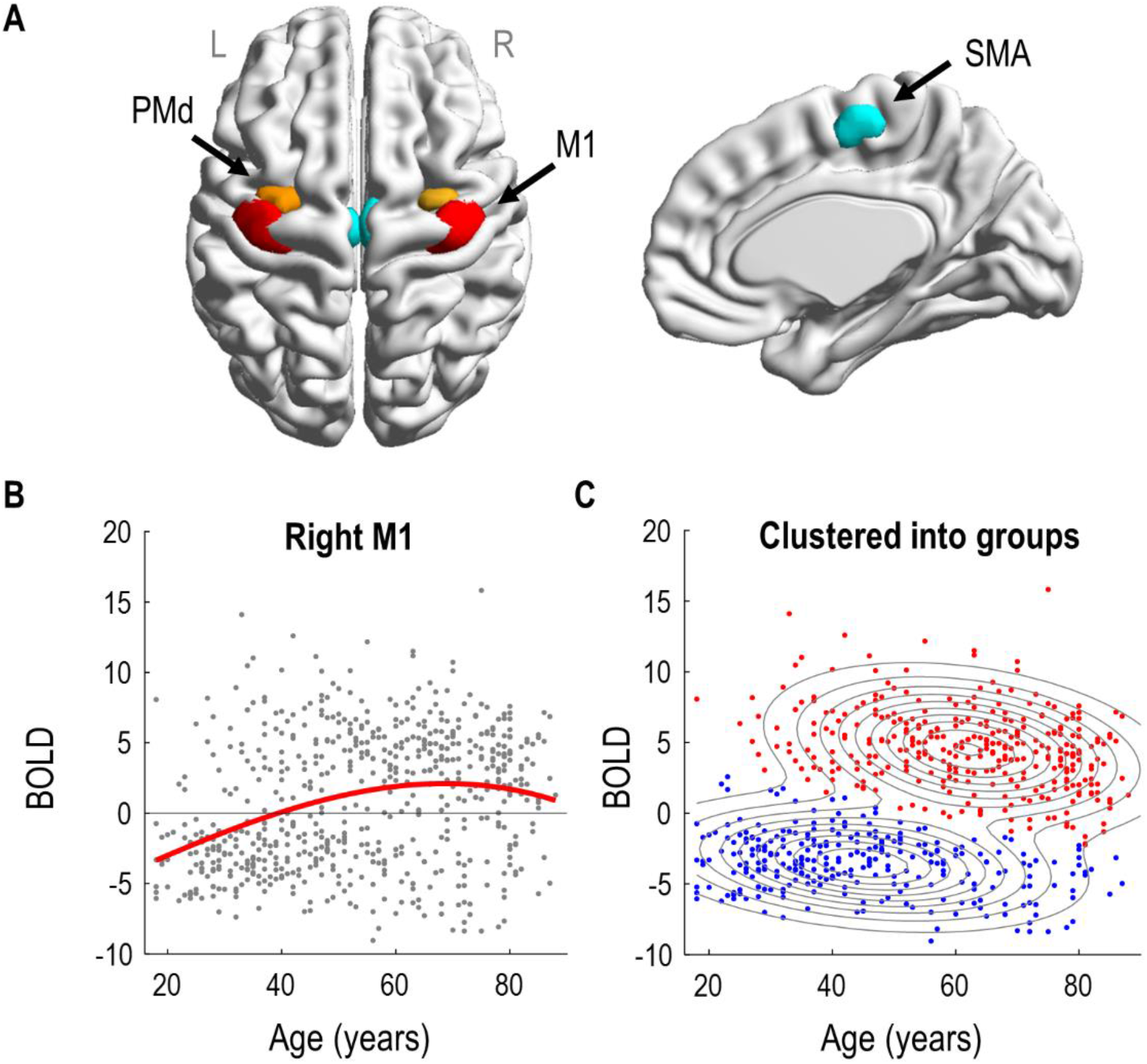
Regions of interest and the effect of age on right M1. **A.** The six regions of interest are shown overlaid on a dorsal view of the cortical surface (left) and on a sagittal view of the right hemisphere (right). M1 = primary motor cortex (red), PMd = dorsal pre-motor cortex (orange), SMA = supplementary motor area (turquoise). **B**. Confound-corrected relationship between age and the estimated neural response to Auditory+Visual trials in rM1, from an initial mass-univariate (SPM) analysis. The red line is the best fit of a third order polynomial, matching the SPM analysis used to identify rM1. **C.** The same rM1 responses with assignments to two clusters using a Gaussian mixture model. Contours indicate the two Gaussians with colours of the individual subjects (dots) indicating cluster assignment. The younger group (with mainly negative BOLD) were described by a 2D-Gaussian with means [44.77 - 3.53] for age and BOLD response respectively, with variances [289.71 5.20] and covariance −11.16. The older group (with mainly positive BOLD) were described by a 2D-Gaussian with means [62.07 4.51], with variances [238.31 8.12] and covariance −12.43. Renderings were performed using BrainNet Viewer (http://www.nitrc.org/projects/bnv/) [40].

The effect of age on right M1 is illustrated in Figure 1B. Each subject’s response is shown as a grey dot, and overlaid upon these is a third order polynomial model fit, as was used to identify the regions in the SPM analysis. (Similar curves for all regions of interest are provided in Figure S3). Visual inspection suggested at least two types of subject. People with negative BOLD responses were more likely to be under 50, whereas people with positive BOLD responses were more likely to be over 50.

To formalise this observation, we fitted a Gaussian mixture model to the same two-dimensional data and used it to cluster the subjects into two groups (Figure 1C). The model was initialized with two clusters, one with subjects who had positive rM1 responses, and the other with subjects who had negative rM1 responses. Following model estimation using the EM algorithm, 46% of subjects were assigned to the group with mainly negative rM1 responses and they were generally younger in age (mean age 48 years, SD 17). The remaining 54% of subjects, with generally positive BOLD responses in rM1, were typically older (mean age 62 years, SD 15). We will refer to these two groups as the negative and positive responders respectively. We compared the log evidence for this two-cluster mixture model, approximated by the AIC and BIC, against the log evidence of a control model with only a single cluster. The two-cluster model was better in both cases (ΔAIC: 173.71, ΔBIC: 146.99). In summary, clustering the subjects into two groups served to collapse the two highly correlated variables of age and rM1 BOLD response into one principle mode of variation, simplifying the connectivity analyses that followed.

Next, we identified the neural dynamics and haemodynamics of the six brain regions shown in Figure 1A that could distinguish negative and positive responders.

### 3.5 First level DCM analysis

Our first connectivity question was whether the multivariate fMRI data from the six regions of interest were best explained as being driven by the left or right hemisphere (SMA and PMd). We formalised each of these two hypotheses as a connectivity model (DCM), as illustrated in Figure 2.

**Figure 2.**
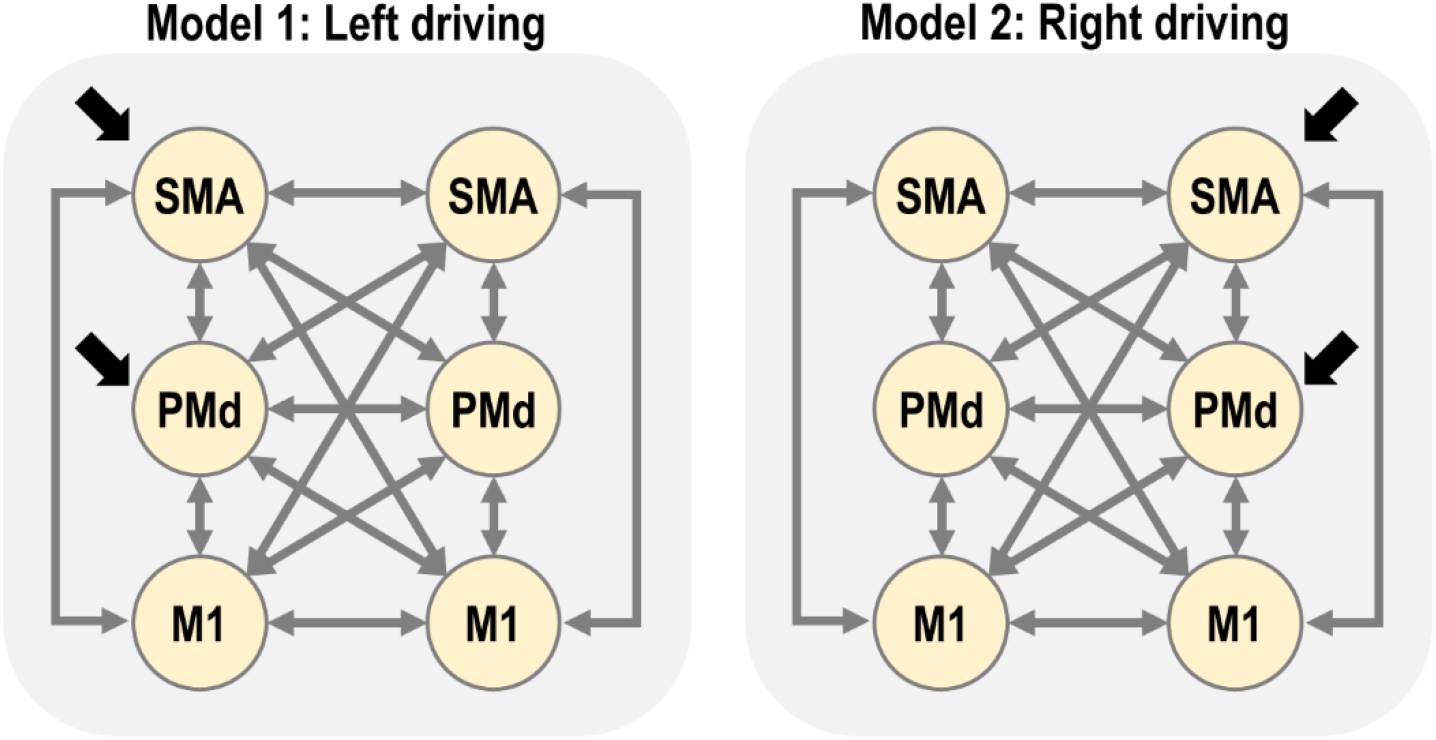
Candidate network architectures. The lighter arrows indicate connections among regions (matrix ***A*** of the DCM neural model, Eq. 1) and the darker arrows indicate the location of driving inputs (matrix ***C*** of the DCM neural model). Self-connections on each region are present but not shown. The left hemisphere of the brain is on the left of each diagram (and in all subsequent figures). SMA=supplementary motor area, PMd=dorsal premotor cortex, M1=primary motor cortex.

After fitting the two DCMs per subject, we checked their explained variance, which serves as a diagnostic for model fitting. More than 10% of variance explained is generally considered acceptable for DCM for fMRI in the time domain (although note that this statistic cannot be used to compare models, as it does not account for differences in model complexity). For model 1 (left driving), the average variance explained over subjects was: lM1=34%, lPMd=22%, lSMA=24%, rM1=25%, rPMd=21%, rSMA=23%. For model 2 (right driving), the average was: lM1=32%, lPMd=22%, lSMA=24%, rM1=25%, rPMd=21% and rSMA=23%. We were therefore satisfied that the models explained a non-trivial amount of variance overall.

### 3.6 Does left or right hemisphere drive the network?

We extended the subject specific DCMs to the group level using the parametric empirical Bayes (PEB) framework. We specified two hierarchical (PEB) models – one for the left driving DCMs and one for the right driving DCMs. Each PEB model consisted of the subject-specific DCMs at the first level and a GLM at the second level. The design matrix of the GLM included regressors to capture the group average strength of each DCM parameter, the difference between positive/negative responder groups for each DCM parameter, and the residual effects of age and the level of rM1 activation after accounting for group.

We performed Bayesian model comparison to identify which of the two PEB models (left or right driving) better explained the group-level data. The log Bayes factor in favour of the left driving model was 2.584 × 10^4^, equivalent to a posterior probability of 1.00. Thus, there was strong evidence that the left (contralateral) hemisphere drove activity in the network. We therefore focus on the left driving model for the remainder of this paper.

To confirm the DCMs captured the data feature of interest – negative BOLD in right (ipsilateral) M1, we plotted the predicted BOLD response from the DCMs, averaged over subjects in each of the two groups (Figure 3). The DCMs predicted negative BOLD in rM1 for the negative responder group (who were generally younger), and positive BOLD for the positive responder group (who were generally older). The peak of the response in rM1 was earlier in the positive responder group, five seconds post-stimulus, and later in the negative responder group, seven seconds post-stimulus, to the nearest half second. This pattern was similar in rSMA and rPMd. Noticeably, the two groups also differed in the BOLD response in the left (contralateral) hemisphere. Negative responders had overall lower amplitude responses in all left hemisphere regions.

**Figure 3.**
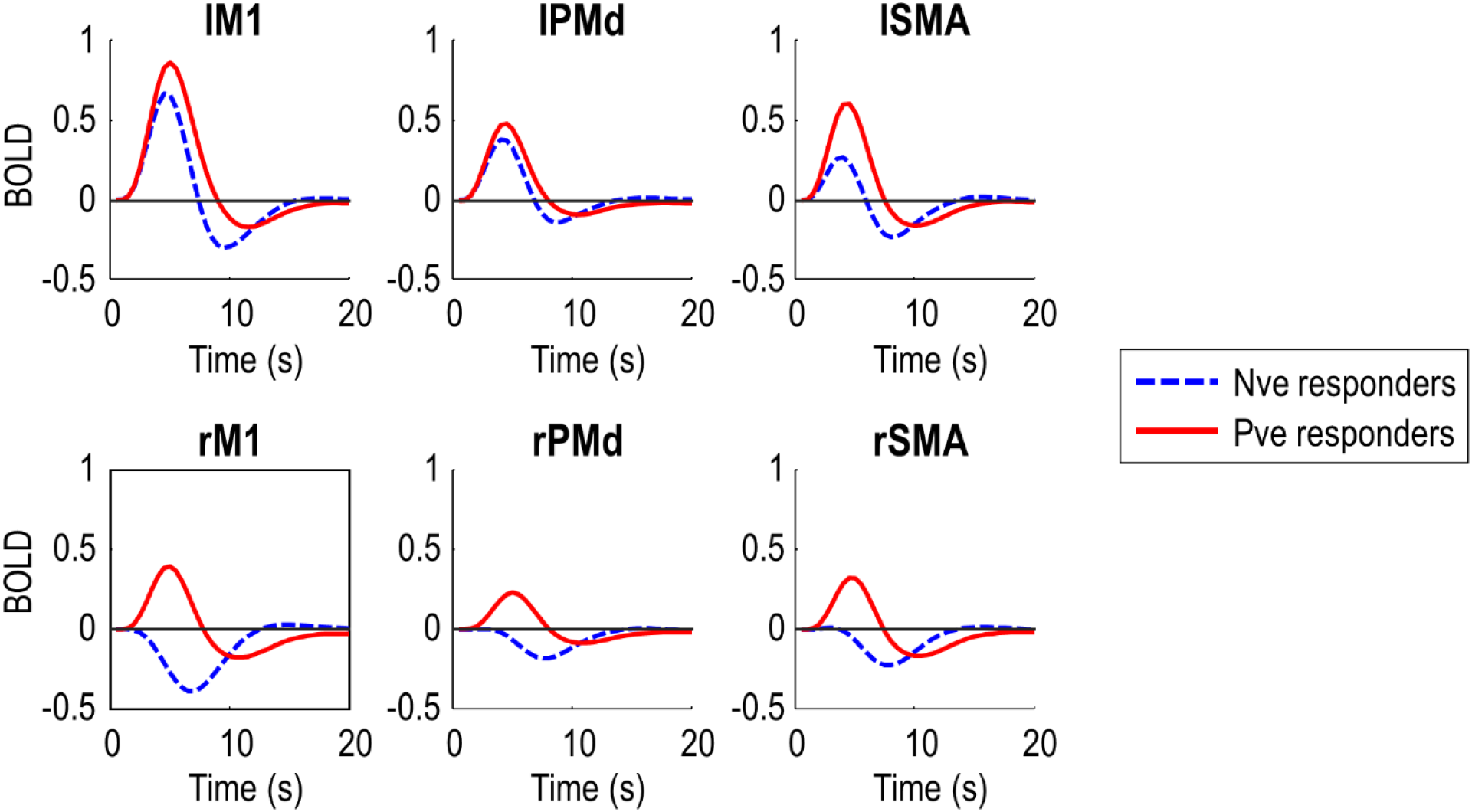
Predicted BOLD responses from the left driving DCM. These responses (first order Volterra kernels) are averaged over subjects from each of the two groups (Figure 1C). The negative (Nve) responder group includes younger subjects, most of whom had negative BOLD responses in rM1. The positive (Pve) responder group includes older subjects, most of whom had positive BOLD responses in rM1. SMA=supplementary motor area, PMd=dorsal premotor cortex, M1=primary motor cortex.

In summary, the DCMs explained a non-trivial amount of variance and were able to capture the data feature of interest – negative BOLD in rM1. Next, we used these models to test specific hypotheses about the causes of the group difference.

### 3.6 Do neural and / or haemodynamic parameters differ between groups?

The PEB model included parameters for the effect of each covariate (average, group, residual age, residual rM1 activation) on each DCM parameter (matrices ***A*** and ***C*** from the neural model, transit and decay from the haemodynamic model). To test whether neural and / or haemodynamic parameters captured the group difference, we compared the evidence for the full hierarchical (PEB) model versus reduced models with particular mixtures of these parameters switched off (fixed at zero).

In more detail, we defined eight sets of parameters: (1) all, (2) A+C, (3) A+transit+decay, (4) C+transit+decay, (5) A only, (6) C only, (7) transit+decay, (8) none. We compared the evidence for 64 candidate PEB models, where each model had its group-average parameters switched on or off according to one of these eight sets, and its parameters encoding the group difference switched on or off according to one of these eight sets. For example, there was a candidate PEB model in which only neural parameters (A+C) were switched on at the group level, with only C parameters differing between groups.

The winning PEB model, with posterior probability 1.00, was the model with all types of parameter (A,C,transit,decay) switched on for both the commonalities across subjects (group average), as well as for the group difference. Thus, both neural and haemodynamic parameters were needed to explain the typical response to the task *and* the differences between groups. We next examined these neural and haemodynamic parameters in turn.

### 3.7 Which neural connections differed between groups?

We identified the specific neural connections that were engaged by the task and those that differed between the two groups. We fitted separate PEB models to the connectivity parameters (matrix ***A***) and driving input parameters (matrix ***C***) from the neural model (Eq. 1). We then performed an automatic search over thousands of reduced PEB models, iteratively pruning mixtures of connections from the model, where doing so did not reduce the free energy. Finally, the parameters from the best models identified during the search were averaged (weighted by the contributing models’ evidence), and are shown in Figure 4.

**Figure 4.**
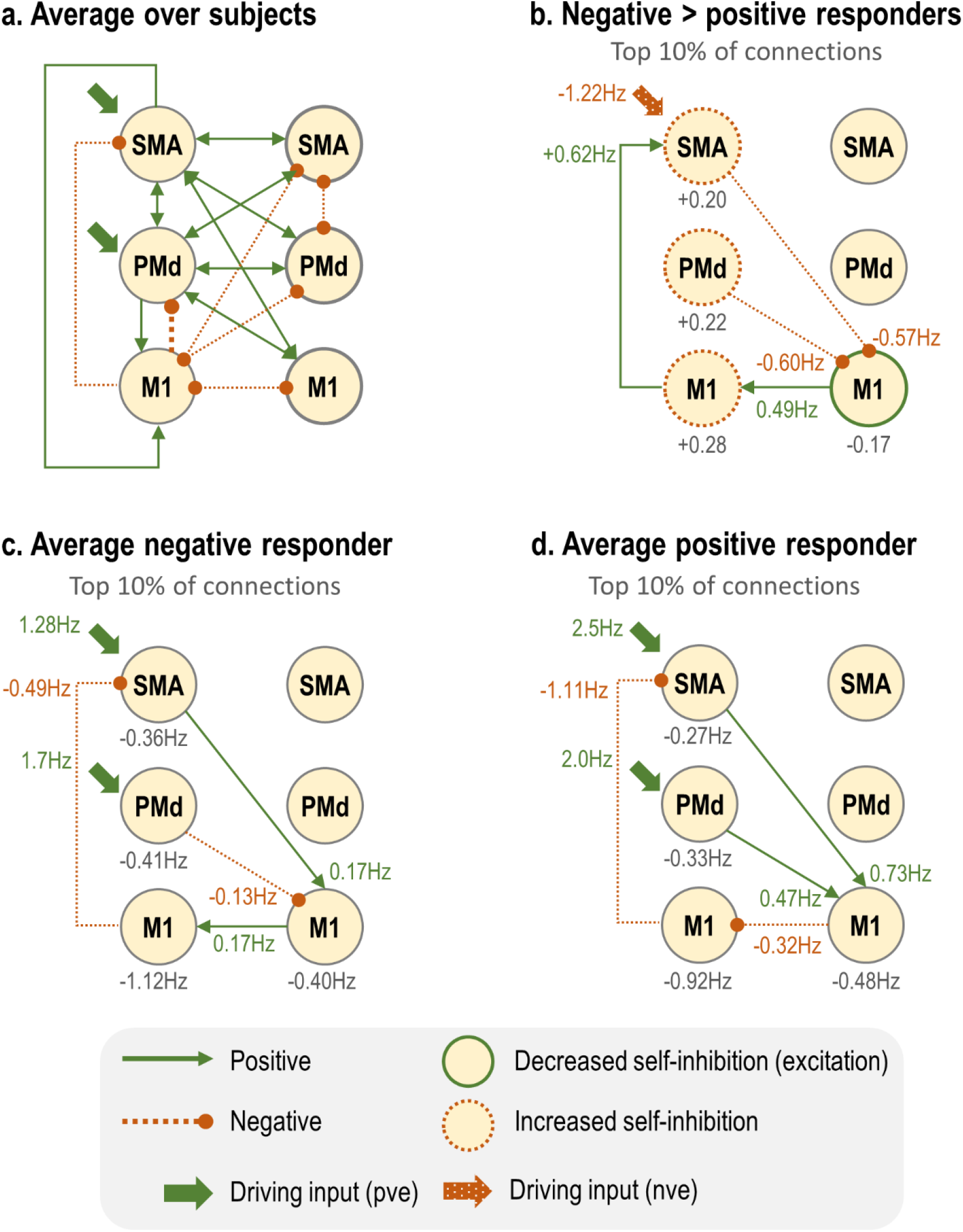
Result of automatic search over neural parameters. A full PEB model was first specified, which contained parameters pertaining to the group average and group difference of all the connections shown in Figure 2 (left). Then an automatic search was conducted to prune away any parameters that did not contribute to the free energy. Finally the parameters of the best models from the search were averaged. The arrows indicate surviving parameters with posterior probability greater than 0.95 (based on the free energy). In panels b-d, the between-region connections are limited to the top 10% by absolute value of the connection strength, for clarity. **(b)** The numbers indicate the difference in connection strengths across groups. Between-region connections, in green and orange, are in units of Hz. Positive values are stronger for the negative responders, negative values are stronger for the positive responders. Light grey numbers indicate the between-groups difference in the log-scaling parameters, which control inhibitory self-connections – positive numbers indicate greater self-inhibition in the negative responders, negative numbers indicate greater self-inhibition in the positive responders. **(c-d)** All parameters displayed in units of Hertz (Hz) for each group separately for clarity. Light grey numbers next to each region indicate self-connections.

Figure 4a shows the parameters encoding the average connectivity across subjects, thresholded at a posterior probability of p > 0.95 (strong evidence) for display purposes. The driving inputs entering SMA and PMd were positive (excitatory), and there was excitatory connectivity among bilateral SMA and PMd. Connections arising from lM1 were inhibitory, and there was reciprocal inhibitory connectivity with rM1. There was also an inhibitory feedback connection: lM1→lSMA.

These average connections contextualise the interesting group differences (Figure 4B). Almost all connections showed group differences (plotted in full in the supplementary materials). We therefore focus on the top 10% of between-region connections, by (absolute) connection strength. For clarity, we also used these values to calculate the group average connectivity for the negative responders (Figure 4c) and the positive responders (Figure 4d). Negative responders had weaker driving input entering lSMA. In turn, they had a more negative inter-hemispheric lSMA→rM1 connection and lPMd→rM1 connection. Conversely, they had a more positive rM1→lM1 connection, and from lM1→lSMA. Some of the self-connections, which control the sensitivity of each region to its inputs, also differed between groups. Negative responders had a small decrease in self-inhibition of rM1, i.e. its responses would be more sustained, and they had increased self-inhibition in all left hemisphere regions, meaning their responses would decay more quickly.

To summarise the neural connectivity parameters, negative responders had less activity in their left hemisphere overall – including weaker driving input to lSMA and greater self-inhibition in all regions. They also had a strong decrease in inter-hemispheric drive from the left hemisphere (lSMA and lPMd) to rM1. Next, we complete our survey of the model parameters by examining those governing haemodynamics.

### 3.8 Which haemodynamic parameters differed between groups?

The model included two kinds of haemodynamic parameters. The *decay* parameter pertains to the neurovascular coupling part of the model. This is a unitless log scaling parameter controlling the rate of decay of the vasoactive signal. The *transit* parameters control the mean transit time of venous blood, i.e., the average time it takes for blood to transverse the venous compartment. Again these are unitless log scaling parameters. There was a single decay parameter pooled across brain regions, whereas transit parameters were estimated on a per-region basis.

We fitted a PEB model to these haemodynamic parameters and applied an automatic search over reduced models to remove any parameters not contributing to the free energy. The parameters of the best models from this search were averaged, and the results are shown in Figure 5. Plots in the top row show the raw estimated log scaling parameters. Within the model, the exponential of these parameters was taken to make them positive in sign, and then they were multiplied by a default value (2 secs for transit, 0.64Hz for decay). To aid interpretation, the plots in the bottom row show the same parameters with this transform applied, for each group of subjects separately.

**Figure 5.**
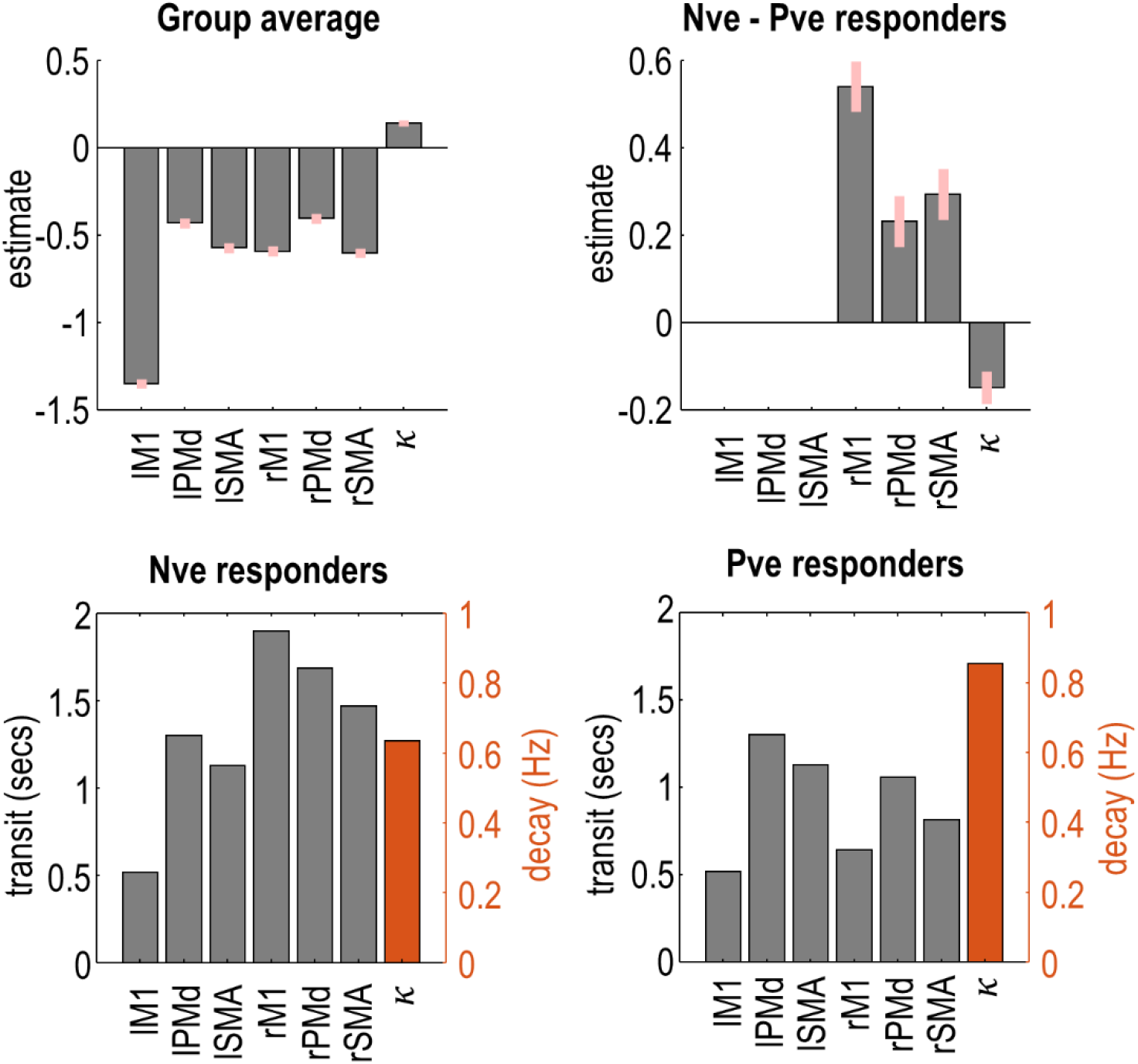
Haemodynamic parameters. **Top row:** The posterior expected values and 90% credible intervals for the haemodynamic log scaling parameters. In each plot, the first six bars are the transit parameters for each region, and the final bar is the decay parameter (which is pooled over regions). **Bottom row:** The posterior expected values, transformed into the units of the underlying model (secs for transit, Hz for decay), plotted separately for each group of subjects.

Negative responders had a longer transit time in all three regions of the right hemisphere, with the largest effect in rM1. Thus, blood flow was more sustained in negative responders, who tended to be younger. There was no difference between groups in left hemisphere regions. Additionally, negative responders had a reduced rate of neurovascular decay (*κ*), i.e., their vasoactive signal was more sustained.

### 3.9 Which parameters explain negative BOLD in rM1?

Having identified model parameters that differed between groups, we next identified which of these parameters determined the sign of the BOLD response in rM1. We specified a DCM that was supplied with the average parameters of subjects in the negative responder group. We then systematically varied each parameter around its estimated value, and recorded the effect on the predicted rM1 BOLD response.

There were only four parameters which, when varied in isolation, were able to switch the virtual subject from a negative to a positive rM1 BOLD response (Figure 6). These were all from the neural, rather than haemodynamic part of the model. Three of these were inter-hemsipheric neural connections: lM1→rM1, lPMd→rM1 and lSMA→rM1. Making any of these connections more positive also made the rM1 BOLD more positive, and delayed the peak of the response. The amplitude of the rM1 BOLD response was most sensitive to input from lM1, then lPMd, and it was least sensitive to lSMA. The fourth parameter was the driving effect of task on lPMd, which had a negative relationship with rM1 BOLD – the more positive the input, the more negative the rM1 BOLD, and vice versa.

**Figure 6.**
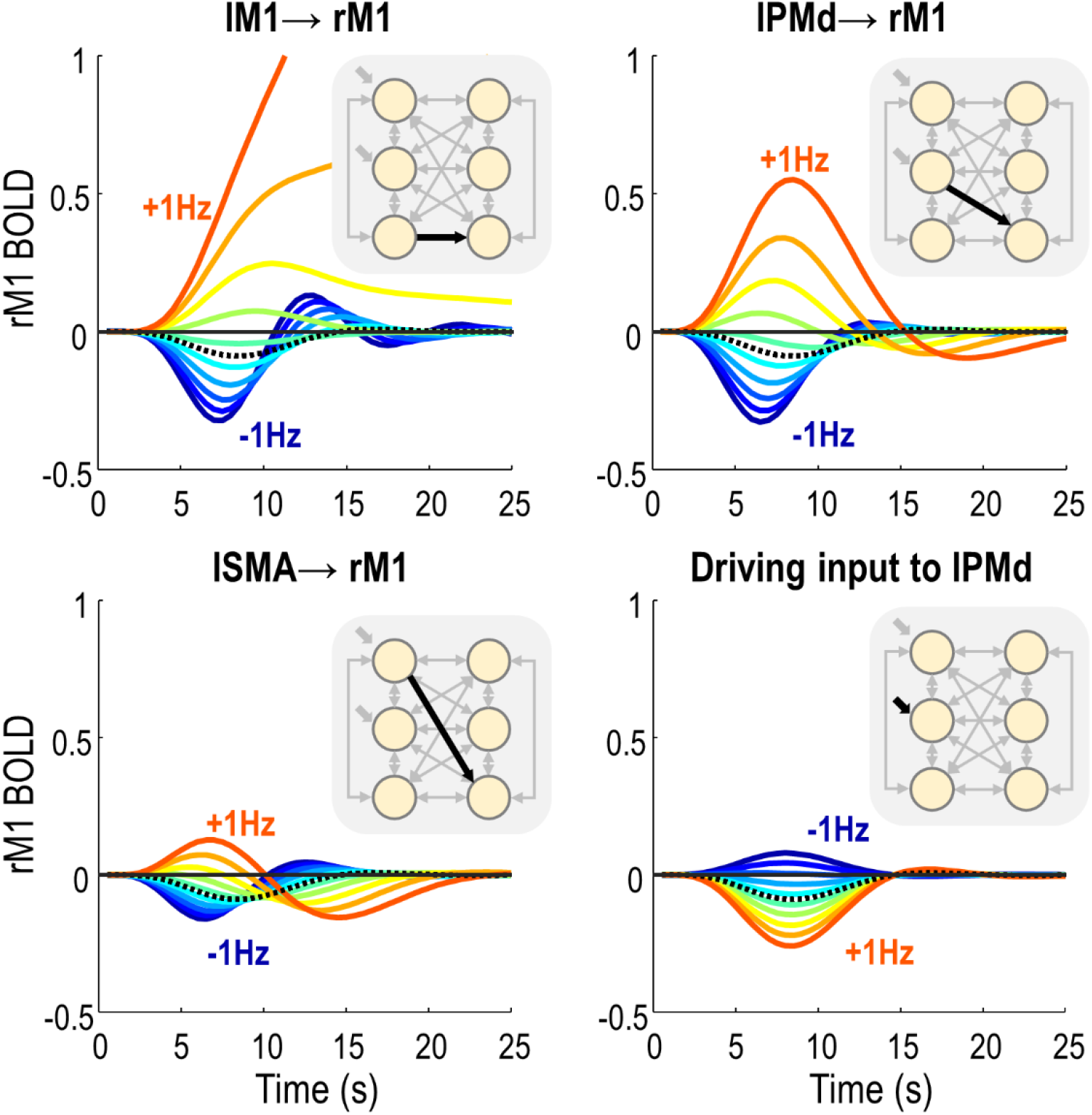
Simulations. Each plot shows the effect of varying one parameter (indicated by the title and the diagram, inset), on the predicted BOLD response in right M1. The black dotted line indicates the predicted BOLD response under the group average parameters for the negative responder group. The coloured lines are simulated BOLD responses, as a consequence of varying each parameter between its estimated value minus 1Hz, to its estimated value plus 1Hz.

Notably, two of these four parameters were those we earlier identified as having large group differences: lPMd→rM1 and lSMA→rM1. Therefore, these two inter-hemispheric connections were sufficient to explain the change in sign in rM1 BOLD between groups, without necessarily needing to modulate the direct lM1→rM1 connection.

To conclude the analysis, we plotted the estimated values for the two inter-hemispheric connections identified above against the amplitude of the response in rM1, as estimated by SPM (Figure 7). Individually, the lSMA→rM1 connection explained 31% of the variance in rM1 response across subjects (R=0.56), and the lPMd→rM1 connection explained 28% (R=0.53). Putting both connections into the same regression model, the overall variance explained in the rM1 response across subjects was 44% (R=0.66).

**Figure 7.**
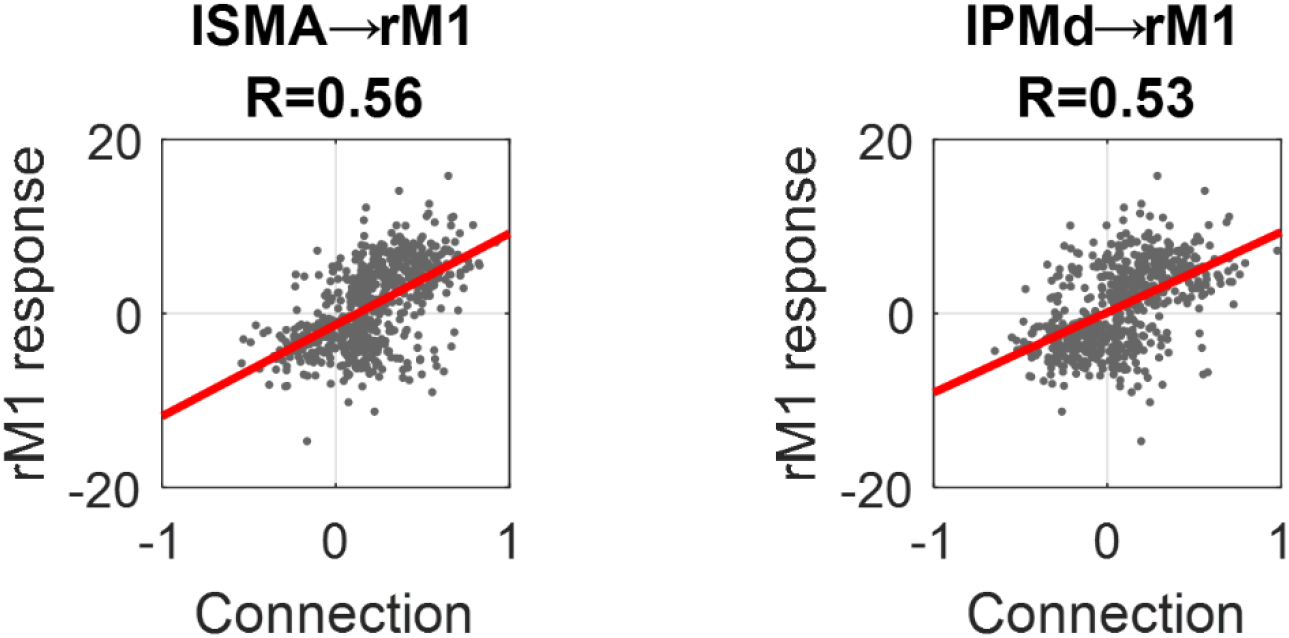
Correlation of inter-hemispheric connections and rM1 response. On each plot, the dots are individual subjects. The horizontal axis is the posterior expected value from the individual subjects’ DCM and the vertical axis is the amplitude of the response in rM1 from the SPM analysis. The line of best fit is shown in red. Pearson’s correlation coefficients are provided in the plot titles.

## 4. Discussion

We investigated why the negative BOLD response (NBR) in ipsilateral M1, elicited by unilateral right-hand button presses, becomes more positive with age. We grouped 635 subjects, aged 18-88, into positive and negative rM1 responders. Using dynamic causal modelling (DCM), we found that only neural (rather than haemodynamic) parameters could explain the positive shift in the rM1 BOLD response with age. In particular, two inter-hemispheric connections, lSMA→rM1 and lPMd→rM1, showed large differences between groups and were sufficient to explain the age-related change from negative to positive BOLD. With these two connections, we were able to explain 44% of the variance in rM1 response across people.

Ageing has a particularly striking effect on the ipsilateral NBR, with only a modest effect on contralateral PBR. Our preliminary SPM analysis, presented in the supplementary materials, confirmed that there was an effect of ageing on the NBR in ipsilateral rM1 and rPMd, as well as bilateral SMA, with no significant effect on contralateral lM1 or lPMd. We were able to capture this difference in BOLD response between hemispheres using dynamic causal models (Figure 3). Ageing therefore does not have a uniform effect on the brain, but rather it alters specific neural populations, brain regions or cognitive processes. This argues against a purely vascular origin for the effects of ageing. To test this and compare the evidence for different underlying mechanisms, we used DCM to partition the variance in subjects’ fMRI data into neural, neurovascular and haemodynamic contributions.

We sought to identify the best model of the fMRI data, where ‘best’ has a precise meaning in a Bayesian setting. Models are compared based on their log evidence, also called the log marginal likelihood, ln *p*(*y*|*m*). This is the probability of having observed the data *y* given the model *m*. This quantity, which is approximated in DCM by the *free energy*, can be decomposed into the accuracy of the model minus its complexity. Therefore, by comparing models of the fMRI data and identifying those with the highest log evidence, we sought to identify models that offered the optimal trade-off between accuracy and simplicity.

We started by specifying two biophysical models (DCMs) for each subject’s data. We opted to use the tried-and-tested deterministic DCM for the fMRI model [41, 42], which made minimal assumptions about neural dynamics, coupled with an established model of haemodynamics [42]. We then extended these models to include between-subjects effects, using the Parametric Empirical Bayes (PEB) framework. Using Bayesian model comparison, we determined that our data were best explained by a network driven by the contralateral, rather than ipsilateral hemisphere. Note that a caveat of this result, and those which follow, is that the fMRI task only involved responses with the right hand. Therefore, we could not separate the left/right hemispheres from the contralateral/ipsilateral hemispheres with these data.

We then used the winning model to investigate which of four types of parameters best explained the difference between groups. We tried different mixtures of the effective connectivity (***A***), the driving inputs (***C***), the rate of decay of neurovascular coupling (*κ*) and / or the haemodynamic time parameters ***τ***. We had hoped that this Bayesian model comparison would whittle down the number of parameters differing between groups. Instead, the group difference was expressed in all four types of parameter. Given the many ways in which ageing affects the brain and its vasculature, this is perhaps not surprising. Furthermore, with such a large number of subjects, we were well powered to detect very small effects. One further technical explanation for why group differences may have been distributed across many parameters stems from the definition of complexity in the free energy. In this context, complexity is a measure of the deviation of the parameters from their prior values, weighted by their prior precisions (the KL-divergence). In the presence of co-varying parameters, moving one parameter far from its prior can have a larger complexity cost than moving many parameters a small distance from their priors [43]. This ‘many-hands-make-light-work’ phenomenon can cause experimental effects to be distributed across many parameters. Given the large number of parameters showing between-groups differences, we focussed on the top 10% of connections by effect size. We additionally examined every parameter using simulations, to determine which of them could determine the sign of the rM1 BOLD response.

One clear pattern that emerged from the estimated parameters was that positive responders, who tended to be older, had stronger neural activity overall, mediated by more positive driving input entering lSMA and more positive inter-hemispheric connections from the contralateral to ipsilateral hemisphere. This gave rise to higher amplitude (positive) BOLD responses in all regions. These results are consistent with many previous studies, which have shown that older participants have stronger activation in response to motor tasks than younger participants [4, 5]. Our results are also largely consistent with two previous DCM studies [26, 27], which found stronger connectivity from the contralateral motor regions to ipsilateral M1 with increasing age. An earlier DCM study identified that SMA inhibits M1 particularly strongly during motor imagery [44], and it would be interesting to determine whether the ageing effects we observed here are specific to motor execution, or if they would also pertain to imagery.

A key advantage of having a generative model of the fMRI data is the ability to perform virtual experiments – lesioning or varying the strength of particular connections - and observing the resulting effects on particular data features. Here, we varied all neural and haemodynamic parameters and recorded the effect on the ipsilateral rM1 BOLD response. We found that only neural, not haemodynamic, parameters were able shift the sign of the BOLD response from negative to positive, and two of these connections also showed substantial between-groups differences: lSMA→rM1 and lPMd→rM1. Increasing either of these connections was sufficient to progress from a negative to positive rM1 BOLD response, in a similar manner to the process of ageing. This was not the case for the lM1→rM1 connection, which speaks to tractography results showing that inter-hemispheric SMA connectivity is far stronger than inter-hemispheric M1 connectivity [35].

Why might these connections become stronger with age? One possibility is that they compensate for age-related decline [45, 46], which could account for the decrease in functional asymmetry across hemispheres (‘HAROLD’). However, this compensation story has recently been challenged. Using the same Cam-CAN dataset as we used here, Knights et al. (2021) found that the level of ipsilateral activity was not associated with behavioural performance, as may be expected if there was a need for compensation that was not entirely fulfilled. Furthermore, including the ipsilateral hemisphere motor activity in a statistical model provided no additional information about the movement being performed than only including the contralateral neural activity [12]. A similar result was found in a study investigating compensation as an explanation for the shift from posterior to anterior regions with age (‘PASA’) [47]. One caveat with those studies, and our study here, is that the tasks used are very simple. It is possible that ipsilateral cortex would encode distinct information if there were more complex movements, particularly if greater demands were made on timing or bilateral coordination.

There may be other physiological, rather than cognitive, explanations for the change in neural inter-hemispheric connectivity, and the resulting de-differentiation of the hemispheres. As set out in the introduction, an age-related reduction in the thickness of the corpus callosum is often noted [21], with associated reduction in IHI, which may alter the dynamics of inter-hemispheric communication. Another candidate mechanism for age-related reduction in brain asymmetry comes from recent work in epigenetics [48]. That study found higher DNA (CpH) methylation in the left hemisphere of the brain than the right, a marker that correlates with the repression of enhancers and promoters in neurons. This asymmetry decreased with age, with the epigenome of the right hemisphere becoming more like the left over the years. This may give rise to (as yet unknown) changes in the neurons of the right hemisphere, reducing hemispheric specialisation.

Neural explanations such as these are consistent with our finding that only neural connectivity parameters, rather than neurovascular or haemodynamic parameters, could account for the shift in ipsilateral BOLD response with age. This also aligns with previous studies that have found a tight coupling between negative BOLD and decreases in neural firing relative to spontaneous activity, in humans [8, 49] and in non-human primates [50]. Additionally, using the Cam-CAN dataset, Tsvetanov et al. found that age-related increases ipsilateral motor responses could not be explained by vascular reactivity, as approximated using the amplitude of resting state fMRI [25]. Nevertheless, there are limitations to our study that may have reduced our ability to detect effects that were specifically neurovascular or haemodynamic. First, we only used one type of data - BOLD fMRI – with no independent measures of neural activity (EEG/MEG) or blood flow (ASL fMRI). We instead separated out neural and non-neural effects using a generative model, which had both region-specific parameters and neurovascular coupling parameters shared across regions. This complemented an efficient experimental design that sampled a wide range of inter-stimulus intervals, enabling the characterisation of non-linearities in the haemodynamic response. Nevertheless, to confidently distinguish vascular effects from neural responses would require the integration of multiple modalities of neuroimaging data. The Cam-CAN dataset includes direct neural recordings - magnetoencephalography (MEG) - of the same subjects who underwent fMRI while performing the motor tasks. These data could be integrated with the fMRI data via a ‘Bayesian fusion’ approach, whereby a common generative model gives rise to both kinds of data [51, 52]. This would enable the evidence to be formally assessed for between-subjects effects having neuronal and / or haemodynamic origins. We plan to apply these methods in future.

A second limitation of our study pertains to the generative model we used. It enabled us to estimate the strength of neurovascular coupling on a per-subject level, however it could not account for excitatory and inhibitory neural populations within each brain region having distinct neurovascular coupling. This may be relevant here, given findings of differential metabolic demand for positive relative to negative BOLD responses [8]. This limitation was first addressed in an extension to DCM by Havlicek et al. [53], who re-parameterised the neurovascular coupling model in tandem with the use of a two-state-per-region neural model. A further step up in biological detail would be to apply a laminar haemodynamic model [54], which together with high-resolution fMRI data (likely at 7T or above) and a laminar model of neural dynamics, could enable a more spatially fine-grained investigation into the genesis of the NBR and its age-related changes.

While the connections we identified could account for a significant amount of the inter-subject variability in rM1 response, there remains a lot of variance to explain. It is clear from the data (Figure 1B) that some 30-year-olds had rM1 responses similar to those of 80-year-olds, while some 80-year-olds had responses similar to the youngest participants in the dataset. Our long-term goal is to understand the reason for these inter-subject differences, and determine whether they correlate with health and clinical measures. We anticipate this will be aided by having a sensitive characterisation of the physiological processes causing the fMRI data, such as the estimates of connection strengths established here. Accompanying this paper we provide the connectivity estimates from each subject, as well as the scripts needed to reproduce the analyses. Hypotheses can then be tested by correlating these model parameters against selected variables from the Cam-CAN dataset, which includes hundreds of lifestyle variables, demographic data as well as physiological measures.

## 5. Conclusions

The novelty of this study was the use of generative models (DCM for fMRI), with parameters fitted to an exceptionally rich fMRI dataset, in order to quantify the neural, neurovascular and haemodynamic contributions to the negative ipsilateral BOLD response across the lifespan. With a simple button pressing task conducted with the right hand, we found that the negative BOLD response in ipsilateral rM1, but not contralateral lM1, became more positive with age. People with positive rM1 BOLD has stronger inter-hemispheric effective connectivity from lSMA and lPMd to rM1, and these two connections were able to explain 44% of the variance in the rM1 response across people. We therefore conclude that inter-hemispheric neural connectivity from lSMA and lPMd are major contributors to the age-related change in the sign of the rM1 BOLD response.

## Supporting information

Supplementary methods and results

## Author Contributions

Conceptualization, P.Z.; investigation, Y.W.T and P.Z.; data curation, E.K. and R.H.; writing— original draft preparation, Y.W.T. and P.Z.; writing—review and editing, P.Z., Y.W.T., E.K. and R.H.; supervision, P.Z. All authors have read and agreed to the published version of the manuscript.

## Institutional Review Board Statement

Data used for this study were previously collected by Cam-CAN and made openly available to researchers. These data were collected in compliance with the Helsinki Declaration and approved by the Cambridgeshire 2 (now East of England-Cambridge Central) Research Ethics Committee (reference: 10/H0308/50).

## Informed Consent Statement

Informed consent was obtained from all subjects involved in the study.

## Data Availability Statement

The data are available on request from this website: https://camcan-archive.mrc-cbu.cam.ac.uk/dataaccess/. Scripts and models will be made publically available following peer review.

## Acknowledgments

Thank you to Dr Edda Bilek for advice on figure preparation.

