## Supplementary methods and results for "Ageing and the ipsilateral M1 BOLD response: a connectivity study"

### 1. Supplementary Methods

#### *1.1 Preparation of behavioural covariates*

We included the 17 covariates for our SPM analyses, listed in Table S1. The age regressor was mean-centred before calculating the higher order terms, and the three age terms were orthogonalised using recursive Gram-Schmidt orthogonalisation (with the Matlab function spm_orth.m).

**Table S1.** Covariates used in the behavioural analysis and their definition.

| | Variable | Cam-CAN home interview code |
|---|---|---|
| 1 | Age | |
| 2 | Age squared | |
| 3 | Age cubed | |
| 4 | Sex | |
| 5 | Handedness [A] | |
| 6 | MR coil | |
| 7 | Smoking [B] | v241, v252, v259 |
| 8 | Blood pressure | v349 |
| 9 | Cholesterol | v355 |
| 10 | Cardiac [C] | v357, v359, v361, v370, v372 |
| 11 | Migraine | v366 |
| 12 | Diabetes | v377 |
| 13 | Height | |
| 14 | Weight | |
| 15 | Blood pressure (systolic) [D] | |
| 16 | Blood pressure (diastolic) [D] | |
| 17 | Resting pulse rate [D] | |
| 18 | Hearing loss | v336, v342 |
| 19 | Hearing test score (left) [E] | v346 |
| 20 | Hearing test score (right) [E] | v344 |
| 21 | Hearing test unavailable | v344, v346 |

(A) Edinburgh handedness inventory (EHI). (B) Whether the subject smokes now, or had previously smoked on at least 100 occasions, or if anyone in their household smokes. (C) Any one of: angina, heart attack, cardiac arrhythmia, pulmonary embolism, deep vein thrombosis. (D) Resting values averaged over the second and third of three measurements per subject. (E) Number of 1Khz tones heard (max 3).

#### *1.2 SPM analysis*

For each subject we specified a general linear model (GLM) with three conditions of interest: auditory+visual trials (collapsed over auditory frequency), auditory only trials and visual only trials. Each condition was modelled as a separate box-car regressor with trials of duration 300ms, which were convolved with the default canonical haemodynamic response function (HRF) in SPM. We also included six regressors encoding head movement and a regressor to model the session mean.

To test for commonalities across subjects and differences due to age, we ran one-sample t-tests across subjects using SPM, separately for the auditory+visual, auditory and visual conditions. The variables in Table S1 were included as covariates. For each of the three models, we computed an F-contrast to test for any effect of age (specified as an identity matrix of dimension three across age, age squared and age cubed, to capture both linear and non-linear effects). Results were rendered onto 3D cortical surfaces using BrainNet Viewer (http://www.nitrc.org/projects/bnv/) [1].

### 2. Supplementary Results

#### *2.1 SPM: average response to auditory+visual trials*

We first identified regions of the brain responding to the task, averaged across all subjects (Figure S1). Most voxels showed task-related effects, which is unsurprising given the large number of subjects and the use of classical statistics. As expected, a particular difference can be seen between hemispheres in sensorimotor cortex, in which the right hemisphere (circled) has negative responses, whereas the left hemispheres has positive responses.

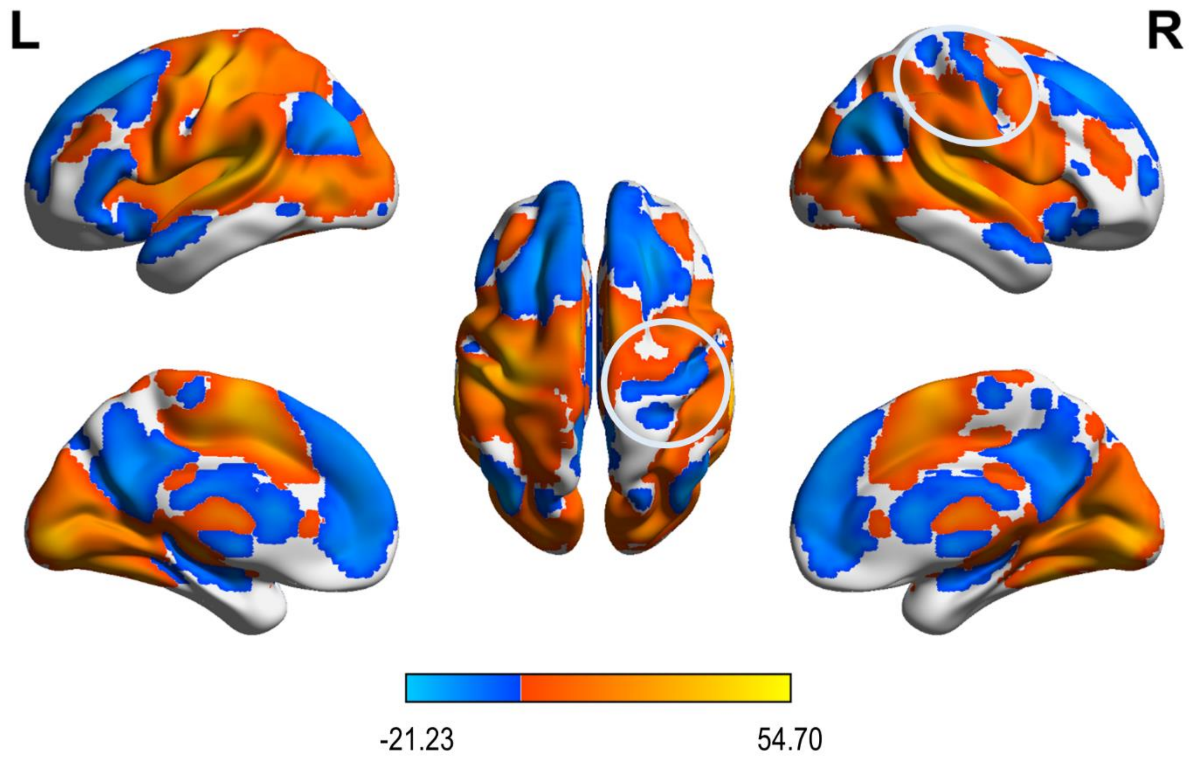


**Figure S1.** The main effect of task (auditory+visual trials). Results of a T-contrast are projected onto the ICBM152 (smoothed) template, thresholded at P < 0.05 family-wise error corrected. Hot colours indicate positive responses (relative to unmodelled time, particularly inter-trial intervals) and cold colours indicate negative responses. The circles highlight negative BOLD in right sensorimotor cortex.

*2.2 SPM: effects of age*

We next identified locations in the brain showing systematic effects of age. We fitted a second level regression model for auditory+visual trials, including age effects as covariates while controlling for other variables that could affect neurovascular coupling (listed in Table S1). An F-contrast for effects of age, covering linear, quadratic and third order terms, identified significant clusters including right sensorimotor cortex, bilateral supplementary motor area (SMA) and bilateral primary auditory cortex (Figure S2 and Table S2). Post-hoc t-tests (not shown) demonstrated that the age effects in bilateral SMA and right sensorimotor cortex were positive (i.e., more positive activity in older people) whereas effects in primary auditory cortex were negative (i.e., less positive activity in older people).

We used an atlas called the Human Motor Area Template (HMAT) [2] to label the regions that constituted the right sensorimotor cluster (arrows in Figure S2). 60% of voxels were in right M1, 23% were in right S1 and 12% were right dorsal premotor cortex. The remaining 5% were outside the atlas.

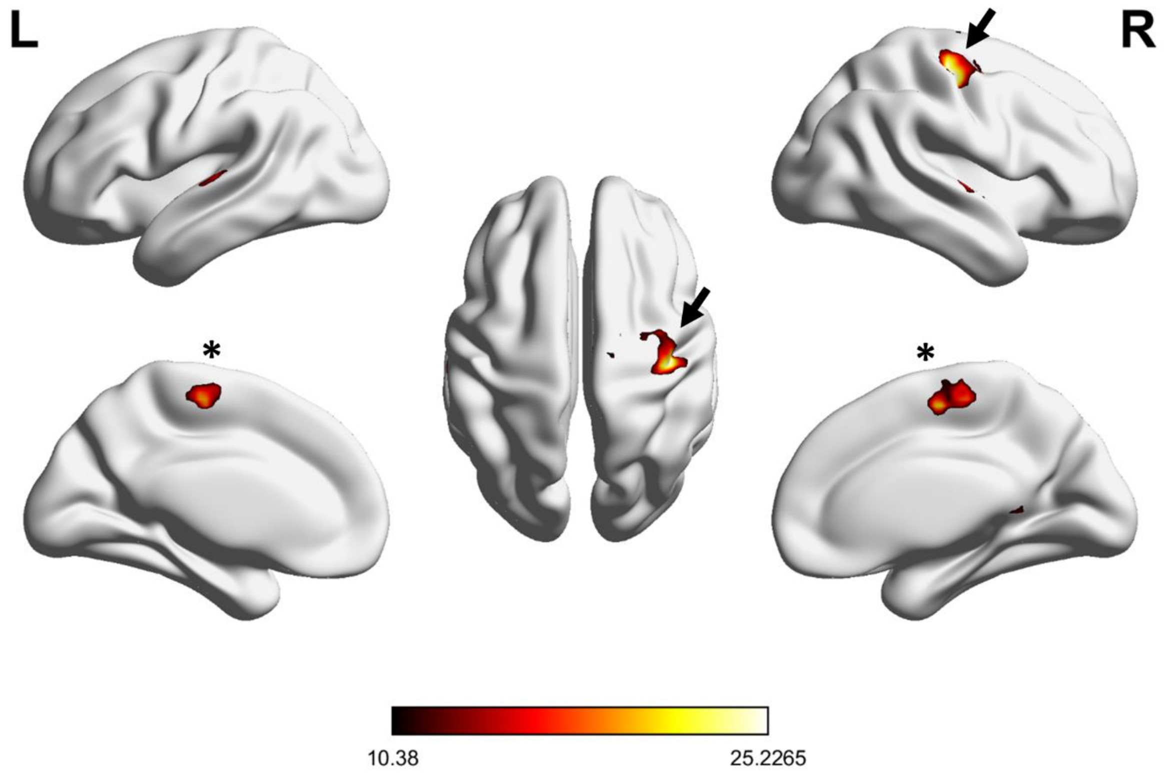


**Figure S2.** Effects of age on the response to auditory+visual trials. Results of an F-contrast are projected onto the ICBM152 (smoothed) template, thresholded at P < 0.05 family-wise error corrected. Arrows indicate a cluster that includes right primary sensory cortex, right primary motor cortex (M1) and right dorsal premotor cortex (dPM). Asterisks indicate bilateral supplementary motor area (SMA). The colour bar indicates the F-statistic.

**Table S2.** Effects of age on whole-brain fMRI results

| **Description** | **Peak XYZ (MNI, mm)** | **Voxels*** | **Peak F** |
|---|---|---|---|
| Right precentral gyrus / postcentral gyrus (M1) | 39 -21 54 | 293 | 27.3 |
| Bilateral SMA | 6 -12 54 | 124 | 21.8 |
| Left planum temporale / Heschl's gyrus | -57 -24 6 | 89 | 17.2 |
| Right planum polare / central opercular cortex / temporal pole | 48 3 -3 | 73 | 19.7 |
| Left cerebellum (V,V1) | -18 -48 -21 | 21 | 16 |
| Right brain stem | 3 -42 -36 | 18 | 20.1 |
| Right precentral gyrus / superior frontal gyrus | 15 -15 69 | 17 | 13.2 |
| Right retrosplenial cortex | 9 -48 6 | 14 | 15.6 |
| Right precentral / postcentral gyrus | 12 -30 69 | 14 | 16.4 |
| Right cerebellum (V, I-IV) | 15 -45 -18 | 8 | 12.3 |

| | | | |
|---|---|---|---|
| Left postcentral / precentral gyrus | -9 -33 69 | 8 | 14.9 |
| Right thalamus | 9 -21 -3 | 6 | 12.6 |
| Left precuneus | -6 -63 24 | 6 | 13.2 |
| Left angular gyrus / lateral occipital cortex | -48 -57 27 | 5 | 12.7 |

* Table truncated for brevity: only clusters with five or more voxels reported.

*2.3 ROI analyses*

We defined eight regions of interest, as described in the main text. Recall that the lSMA, rSMA, rdPM and rM1 were identified in the SPM analysis on the basis of having a non-linear (3$^{rd}$ order) mapping between response and age. The remaining two regions, ldPM and lM1, were homologues of their right hemisphere counterparts.

We extracted first-level GLM parameters corresponding to the Auditory+Visual condition, averaged over voxels within each region. To confirm that the ROI-averaged results qualitatively matched the voxel-wise results, we calculated (post-hoc) linear correlations between response and age, which unsurprisingly were significant in lSMA, rSMA, rdPM and rM1. To illustrate the non-linear association between response and age, we fitted 3$^{rd}$ order polynomials to the ROI-level results (red curves in Figure S3). A preponderance of negative responses in young subjects is particularly apparent for all right hemisphere regions and left SMA.

Finally, we checked whether any of the confounds we defined (Table S1) had significant effects. We specified an F-contrast as an identity matrix over the confound variables. This was essentially a model comparison of the GLM with versus without the confounds. There were no significant effects across the whole brain. We therefore did not include these confounds in the connectivity analyses that followed.

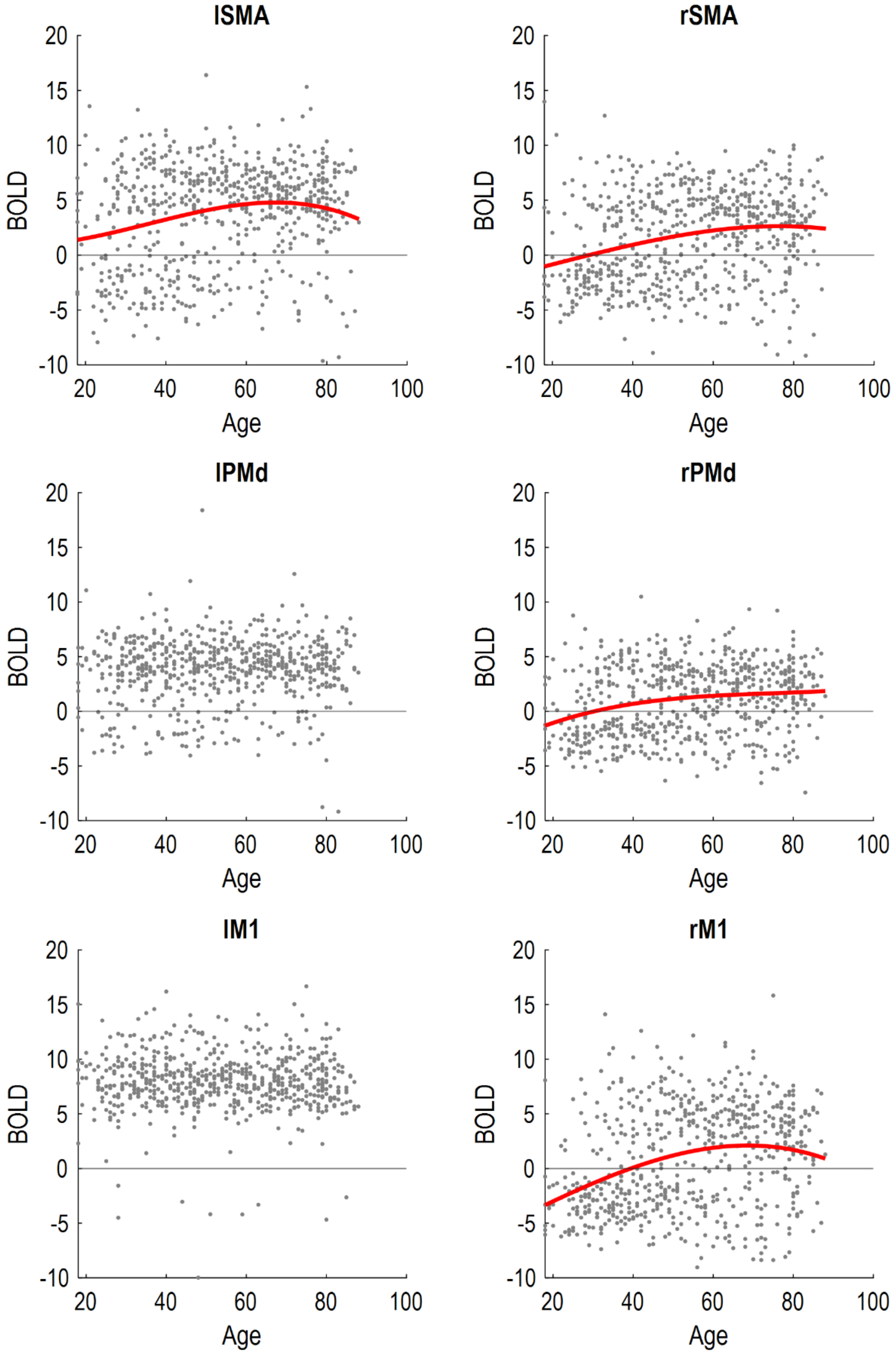


**Figure S3.** Correlation between age and response to auditory+visual trials. Non-linear curves of best fit (red lines) are plotted for those regions with a significant linear correlation (** $p < 0.05$ Bonferroni corrected).

*2.4 Neural parameters*

Figure S4 shows all the neural connectivity parameters from the PEB model, at the group level, following Bayesian model reduction. This procedure pruned any parameters from the model that did not contribute to the log evidence. Strikingly, there was substantially reduced driving input to lSMA (right hand plots) in negative responders, who were generally younger.

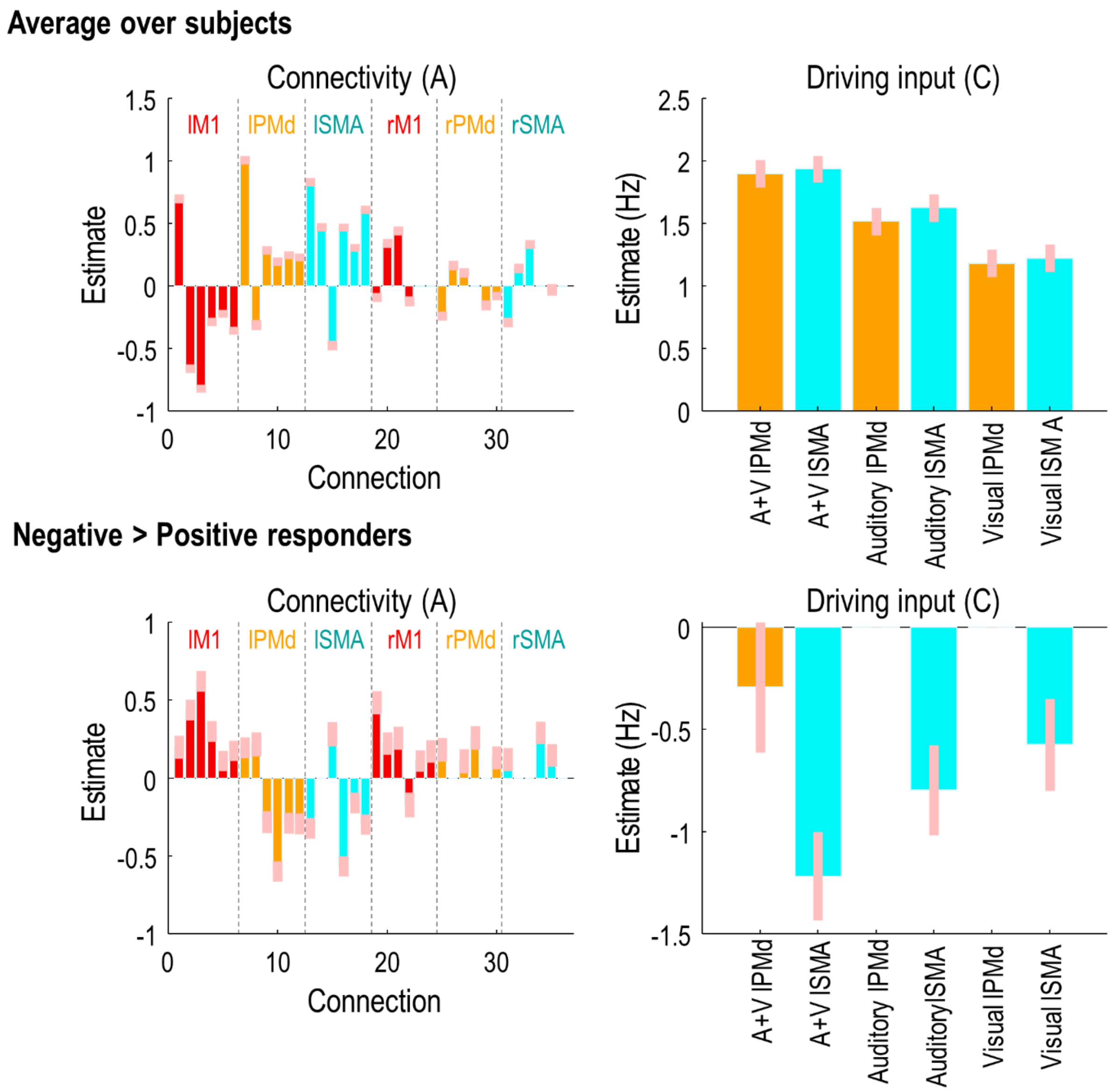


**Figure S4.** Posterior parameter estimates of the neural part of the model. DCM parameters were summarised at the group level using PEB. Parameters that did not contribute to the group-level log evidence (free energy) were pruned using Bayesian model reduction. Colours match Figure 1A of the main text. **Top**: the commonalities across subjects (group average). For the connectivity parameters (left), each group of six bars show the outgoing connections from the labelled region, to: (1) lM1, (2) lPMd, (3) lSMA, (4) rM1, (5) rPMd, (6) rSMA. Pink error bars are 90% Bayesian credible intervals. **Bottom:** The effect of group, where positive values indicate more positive parameter estimates for the younger group (negative responders), whereas negative values indicate more positive values for the older group (positive responders).
